# Can you catch Ebola from a stork bite? Inductive reasoning influences on zoonosis risk perception

**DOI:** 10.1101/084590

**Authors:** Tyler Davis, Micah B. Goldwater, Molly E. Ireland, Nicholas Gaylord, Jason Van Allen

**Author notes:** Corresponding author: Tyler Davis, Psychology Building, MS 2051 Department of Psychological Sciences Lubbock, TX 79409.

## Abstract

Emerging zoonoses are a prominent global health threat. Human beliefs are central to drivers of emerging zoonoses, yet little is known about the factors that influence perceived risks of animal contact. We present an inductive account of zoonosis risk perception, suggesting that beliefs about the range of animals that are able to transmit diseases to each other influence zoonosis risk perception. Consistent with our account, in Study 1, we find that participants who endorse higher likelihoods of cross-species disease transmission have stronger intention to report animal bites. In Study 2, using real world descriptions of Ebola virus from the WHO and CDC, we find that communications conveying a broader range of animals as susceptible increase intentions to report animal bites and decrease perceived safety of wild game meat. These results suggest that cognitive factors may be harnessed to modulate zoonosis risk perception and combat emerging infectious diseases.

## Introduction

Emerging infectious diseases are major economic and public health concerns. A majority of such diseases are of zoonotic origin [1,2]. Drivers of emerging zoonoses include consumption of wild game meat, livestock grazing practices, and adverse contact with animals through bites and handling of carcasses [3,4]. For all of these drivers, human-animal interaction plays a critical role in disease emergence. However, little is known about how people reason about the potential risks in such scenarios [5].

An interdisciplinary approach is necessary to understand how reasoning processes influence the way humans interact with potential zoonosis vectors. One such approach is One Health, which places a major focus on understanding how human-animal interactions impact emerging diseases and developing cost-effective strategies for preventing disease outbreaks before they start [6,7]. Cognitive psychology has the potential to inform research on human-animal interactions, but has been mostly absent from One Health. The present work aims to bridge this gap by examining cognitive principles that influence zoonosis risk perception and how they can be harnessed to shape communications regarding disease risk.

Despite a lack of cognitive research in One Health, several recent studies have examined factors that determine whether people will eat wild game meat [8] and report adverse animal contact, such as bites, to a health professional [9]. One study that is particularly suggestive of underlying cognitive influences found that survey respondents were more likely to report dog bites if they knew bats could transmit rabies to humans [9]. At first glance, these results are surprising–people’s inferences about one species appear to be influenced by their knowledge of a completely different species.

The finding that knowledge about one animal can affect beliefs about other animals may be partly accounted for by two cognitive principles from research on how people make inductive inferences: *premise number* and *premise diversity* [10,11]. According to the premise number principle, people are more confident in inferences that apply to a large number of category members [12,13]. For example, participants told that lions *and* giraffes use norepinephrine as a neurotransmitter will tend to judge rabbits as more likely to use norepinephrine than those told that only lions *or* giraffes do so. In terms of zoonosis risk, knowing that multiple mammals can transmit a disease to humans may increase the perceived likelihood of humans contracting that disease from another animal’s bite. According to the premise diversity principle, people find inferences sound to the extent that they are known to hold for a wider range of category members [14,15]. For example, participants told that *lions and giraffes* use norepinephrine as a neurotransmitter will tend to judge a generalization to rabbits as stronger than participants told that *lions and tigers* use norepinephrine as a neurotransmitter. In terms of zoonosis risk, knowing that both dogs and bats can transmit rabies may increase perceptions of human risk because they are often viewed as very different members of the mammal category.

Although premise number and diversity are plausibly related to the previous observations surrounding bite reporting intentions, research on inductive reasoning has not been extended to work on risk perception in health or real-world decision making about health behaviors. Indeed, basic research on induction often focuses on the underlying cognitive mechanisms by abstracting beyond applications in any particular domain. Moreover, although people’s judgments of infectious disease contagion has been studied in social and health psychology research [16,17], these studies have focused on more affective and cultural influence on transmission beliefs, and not on fine-grained inductive principles related to generalization of contagion beliefs [18].

In the present work, we test two specific hypotheses from our theory that inductive reasoning principles influence zoonosis risk perception. First, consistent with the aforementioned rabies study, individual differences in perceived risk from animal contact should be associated with individual differences in beliefs about interspecies disease transmission. Second, perceptions of risk to humans should increase as a result of communications depicting transmissibility amongst a wider range of species.

## Study 1

The goal of Study 1 was to examine whether intentions to report animal bites are associated with beliefs about interspecies disease transmission. Specifically, based on the premise number principle, we hypothesized that individuals who endorse stronger likelihoods of disease transmission between a number of different animal species would be more likely to perceive human risks from animal bites. To test this hypothesis, we conducted a survey measuring intentions to report bites from a number of common mammals and birds along with judgments of interspecies disease transmission likelihood for a fictitious novel disease.

## Methods

Participants were 289 adults (55% men; mean age = 33.6, *SD* = 10.2) who completed an online survey. The survey was available to Mechanical Turk workers in the following countries where English is the primary language: USA, Australia, Canada, Great Britain, Ireland, New Zealand, and the Bahamas. The majority of participants had undergraduate (48.8%) or advanced degrees (8.7%). The sample was predominantly White (80.6%), with 5.9% Asian, 3.8% Black, 6.9% Hispanic, 1% Native American or Alaskan Native, and 1.7% other ethnicities. A majority of the sample (73.4%) reported currently owning a pet. Participants were compensated $2 for participation in the survey. Informed consent was obtained from all individual participants included in the study. All protocols were approved by the Institutional Review Board of Texas Tech University. All survey materials, analyses, and data are available at http://osf.io/4r79f

### Questionnaire design

The study materials consisted of an electronic survey containing sections on demographics, bite reporting intentions, and species-to-species disease transmission beliefs. Demographics questions included sex, sexual orientation, ethnicity, education level, mother’s and father’s education level, language(s) spoken, and pet ownership. Previous exploratory analyses that we do not report examined whether any of these demographic variables were moderators of our effects.

In the bite reporting section, participants were asked to judge their likelihood of reporting bites from various target animals to a health professional. Participants were told that a health professional could include anything from a doctor or a nurse to a health advice hotline. Participants judged likelihood of reporting for each animal using a slider that could be adjusted in units of 1 from 0-100. The numeric scale also included descriptive labels (*Very Unlikely*, *Unlikely*, *Somewhat Unlikely*, *Undecided*, *Somewhat Likely*, *Likely*, and *Very Likely*) presented above the slider to facilitate consistent use of the scale. Mammal and bird reporting were presented in a random order on separate screens. Mammals included dogs, skunks, monkeys, bats, and squirrels. Birds included grackles, swans, robins, blue jays, and peacocks.

The species-to-species disease transmission beliefs section employed the same sliding scales as the bite reporting section, but participants were asked to rate the likelihood of between-animal disease transmission for a hypothetical new disease. For each question, participants were told, *“*Scientists discover that a new disease can infect the liver tissue of [premise animal]. How likely is it that this disease can infect the following animals: [conclusion animals]?” The conclusion animals were listed on separate lines with individual scales (0-100) after the premise prompt. Premise animals included bats, dogs, skunks, monkeys, grackles, blue jays, swans, peacocks.

Conclusion animals included bats, dogs, skunks, monkeys, squirrels, grackles, robins, blue jays, swans, and peacocks. Fewer premise animals were used than conclusion categories so that less time would be required to complete the survey and to reduce participant attrition. Animals only appeared as conclusion categories when they were not the premise animal. Premises were presented in a random order on separate screens.

## Results

Intentions to report mammal and bird bites were highly reliable within person (mammals: Cronbach’s *α* = 0.86; birds: *α* = 0.95), as were judgments of interspecies disease transmission (mammal-to-mammal: *α* = 0.96; bird-to-bird: *α* = 0.97; between birds and mammals: *α* = 0.99). Nonetheless, linear mixed effects models revealed that intentions to report bites varied considerably between different animal species [Mammals: *F*(4,1152) = 111.1, *p* < .001, *ηp*^2^ = 0.28; Birds: *F*(4,1152) = 35.23, *p* < .001, *η*p^2^ = .11] and ratings of interspecies disease transmission varied between the different premise types [mammal-to-mammal, bird-to-bird, between birds and mammals; *F*(2,576) = 356.3, *p* < .001, *ηp*^2^ = 0.55; see Figs 2 and 3 for individual premise effects]. Intentions to report bites were stronger for mammals than for birds [*t*(288) = 27.06, *p* < .001, *d* = 1.59; Figs 1A,1B], and diseases were rated as more likely to be transmissible within mammals or birds than between mammals and birds [mammal-to-mammal vs between birds and mammals, *t*(288) = 18.77, *p* < .001, *d* = 1.10; bird-to-bird vs between birds and mammals, *t*(288) = 23.42, *p* < .001, *d* = 1.38]. Consistent with previous research suggesting bats are viewed as similar to both mammals and birds [19], bats were rated as more likely to share diseases with birds [*t*(288) = 13.42, *p* < .001, *d* = 0.79] and less likely to share diseases with mammals [*t*(288) = 7.03, *p* < .001, *d* = 0.41] than were other mammals.

**Fig 1.**
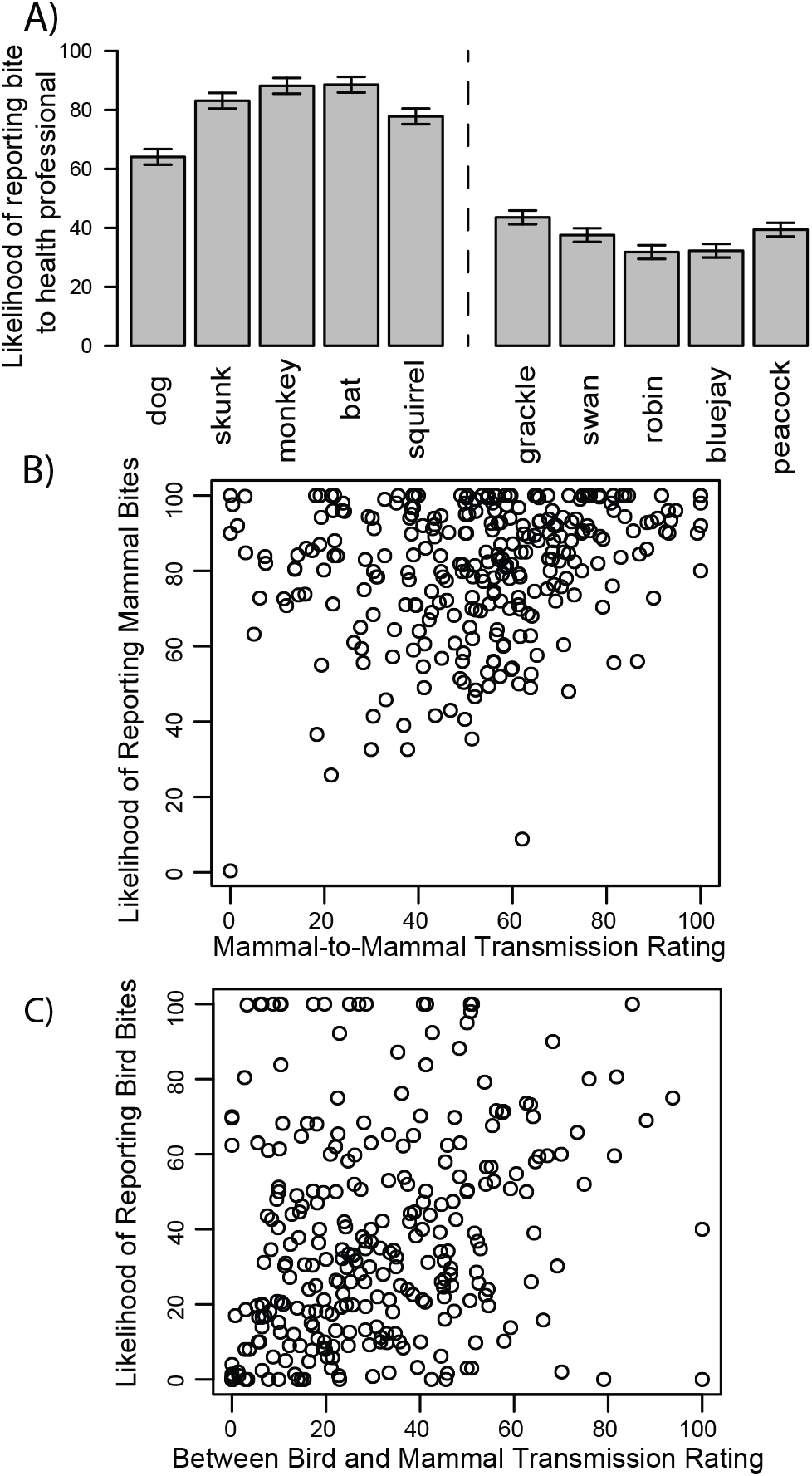
Intentions to report animal bites and associations with transmission ratings. (A) Intentions to report animal bites. (B) Association between intentions to report mammal bites and mammal-to-mammal disease transmission ratings. (C) Association between intentions to report bird bites and between bird and mammal disease transmission ratings. Error bars reflect 95% within-subject confidence intervals.

**Fig 2.**
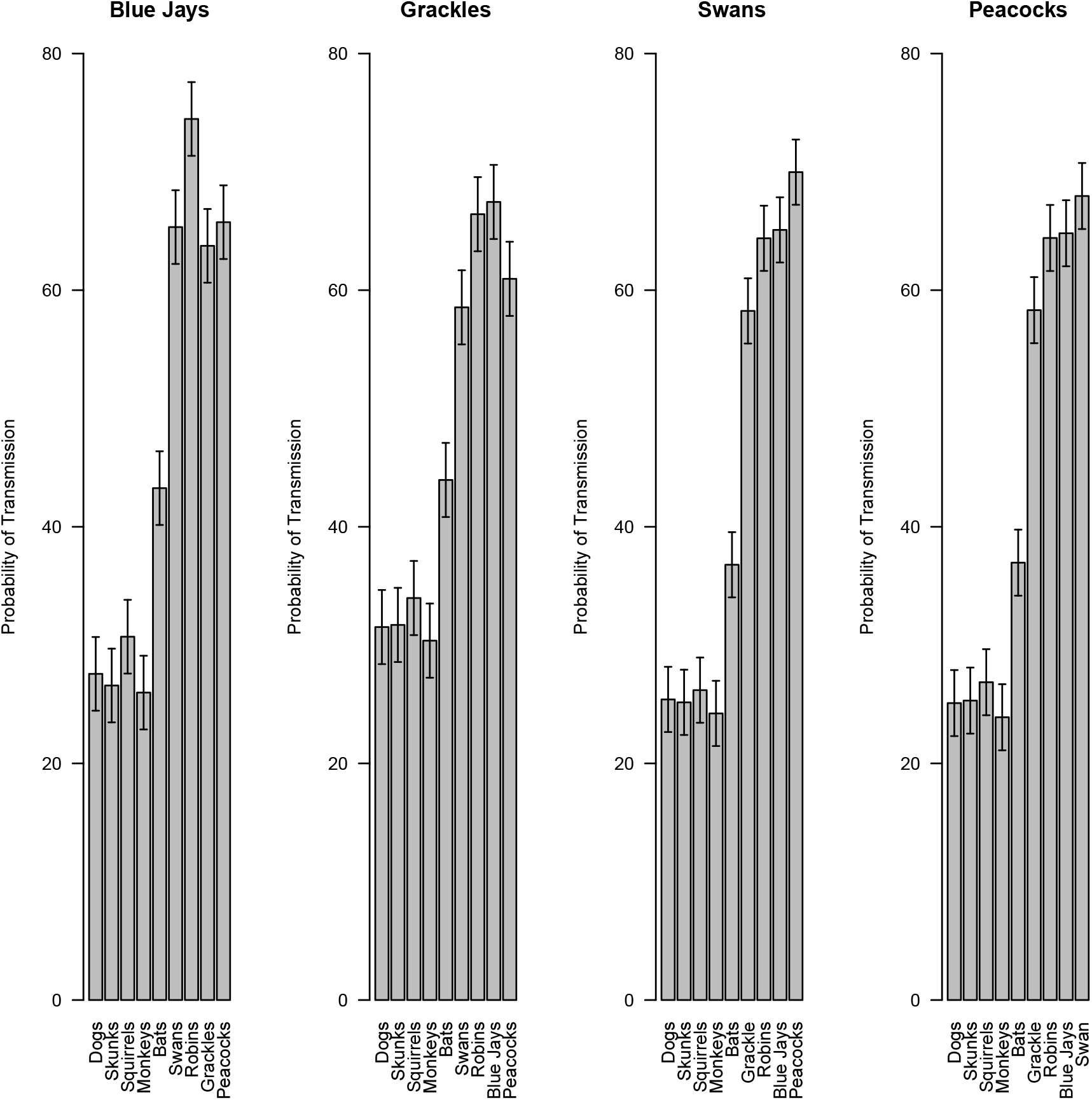
Subjective probabilities of bird disease transmission. Subjective probabilities of disease transmission between each premise animal and conclusion animal for bird premise categories. Graphs are separated by premise category. The bars reflect different conclusion categories. Thus the first bar in the first graph is the mean perceived likelihood of a blue jay disease being transmitted to a dog. Error bars represent 95% confidence intervals calculated from separate linear mixed effects models examining how species-to-species disease transmission beliefs varied for each of the premise birds.

**Fig 3.**
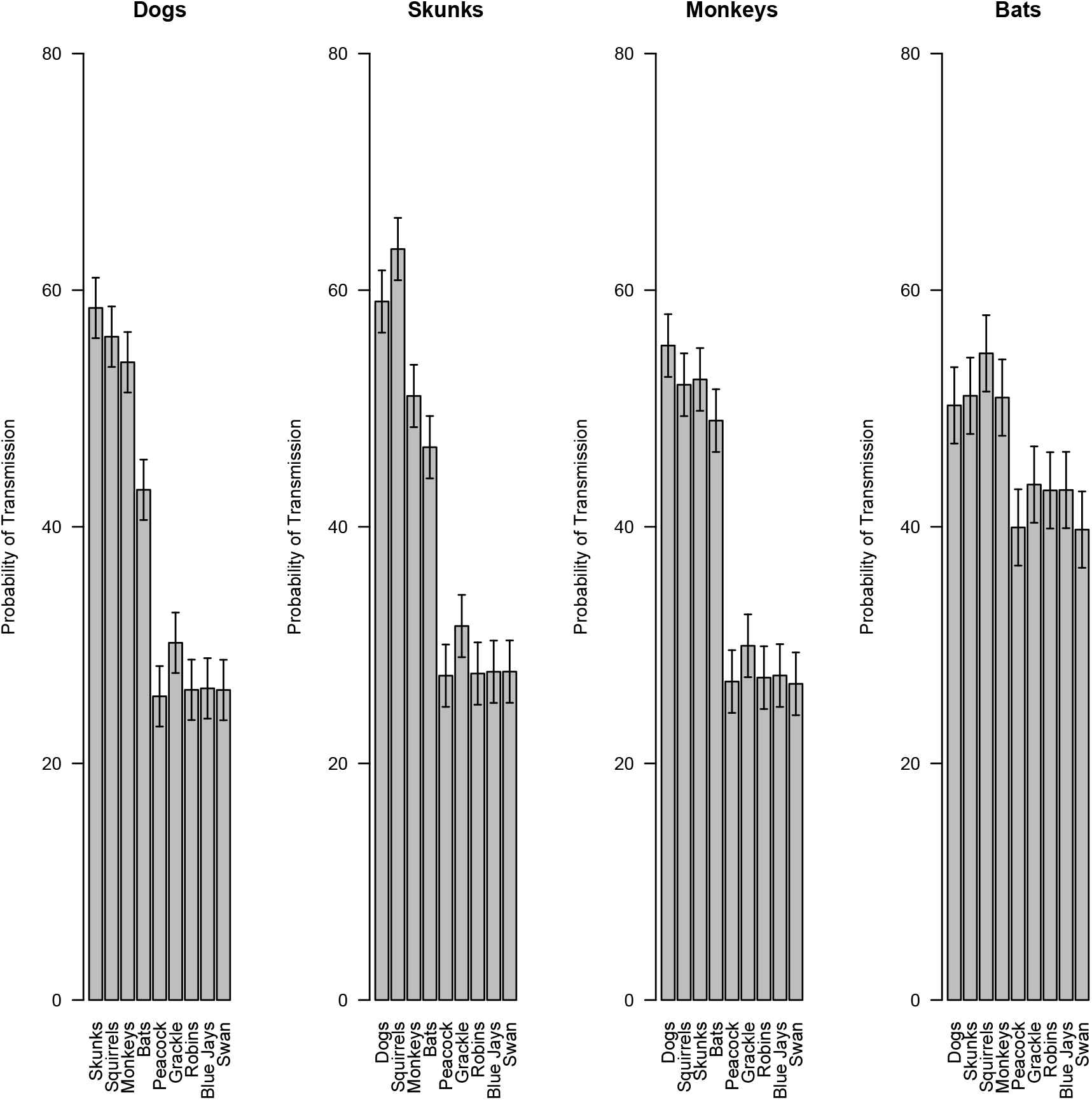
Subjective probabilities of mammal disease transmission. Subjective probabilities of disease transmission between each premise animal and conclusion animal for mammal premise categories. Graphs are separated by premise category. The bars reflect different conclusion categories. Thus the first bar in the first graph is the mean perceived likelihood of a dog disease being transmitted to a squirrel. Error bars represent 95% confidence intervals calculated from separate linear mixed effects models examining how species-to-species disease transmission beliefs varied for each of the premise mammals.

For our primary hypotheses about the association between interspecies disease transmission judgments and bite reporting intentions, we found that individual differences in endorsement of bird-to-bird and mammal-to-mammal disease transmission were both positively associated with individual differences in intentions to report mammal bites [Mammal-to-mammal: Kendall’s τ = .147, *ρ* < .001 (Fig 1B); Bird-to–bird: τ = .140, *ρ* < .001; Between birds and mammals: τ = .009]. Consistent with the premise number principle, endorsing greater odds of interspecies disease transmission was associated with stronger intentions to report mammal bites. For bird bites, only ratings of disease transmission between birds and mammals were associated with reporting intentions [Mammal-to-mammal τ = .043, Bird-to-bird τ = .077, Between birds and mammals τ = .219, *p* < .001 (Fig 1C)]. Coupled with weaker intentions to report bird bites overall, these results suggest that participants may only judge birds as risky (in terms of zoonosis) to the extent they believe birds and mammals can share diseases.

## Discussion

Study 1’s results suggest that inductive reasoning principles underlie people’s perceptions of zoonosis risk. However, because the results are correlational, it is difficult to infer the causal direction between the beliefs about interspecies disease transmission and bite reporting. It is possible, for example, that both interspecies disease transmission and bite reporting ratings are influenced by a common underlying factor, such as individual differences in beliefs about contagion [20] or risk attitudes [21]. Moreover, because the results examine individual differences, it is not clear whether such inductive reason principles could be harnessed to influence people’s beliefs about the risks associated with animal contact.

## Study 2

The goal of Study 2 was to experimentally test whether it is possible to influence people’s perceptions of zoonosis risk through framing communications to convey a greater range of animals as being susceptible to a disease. As a case study, communications about Ebola virus vary in terms of how they describe the range of susceptible animals. The Center for Disease Control’s factsheet [22] lists contact with fruit bats and nonhuman primates (apes and monkeys) as sources of human Ebola infection. Contrastingly, the World Health Organization’s factsheet [23] lists chimpanzees, gorillas, fruit bats, monkeys, forest antelope, and porcupines. According to the premise diversity principle, the WHO’s factsheet should lead to stronger perceptions of Ebola risk from animal contact because it lists a broader range of animals as sources of human Ebola infection. In Study 2, to test whether communications with higher premise diversity would lead to stronger perceptions of risk, we gave participants two different communications about Ebola derived from the WHO and CDC’s factsheets. These communications were tailored from the published factsheets to control all other differences.

## Method

Participants were 152 adults (49% men; mean age = 34.0, *SD* = 10.2) who completed an online survey. Most participants had either undergraduate (49.3%) or advanced degrees (8.6%). The sample was predominantly White (79.6%), with 9.9% Asian, 5.9% Black, 1.3% Hispanic, 0.7% Native American or Alaskan Native, and 0.7% other ethnicities. A majority of the sample (84.2%) reported currently owning a pet and 94.7% reported eating meat. All protocols were approved by the Institutional Review Board of Texas Tech University. All survey materials, analyses, and data are available at http://osf.io/4r79f

### Experimental design

The study materials consisted of an electronic survey containing a demographics section, an experimentally manipulated reading prompt about Ebola, derived from the online CDC and WHO factsheets, an Ebola susceptibility section, a bite reporting intentions section, and a meat safety section. Demographics questions were the same as in Study 1 except for additional questions about meat consumption. Participants reporting that they eat meat were asked additional questions on how often they eat meat, how often they would like to eat meat, and how often they eat wild game meat. If participants answered that they wanted to eat a different amount of meat than they currently do, they were further asked to rate how much personal appearance, ethics, environment, health, spouse’s desires, cost, availability, and taste impacted this discrepancy.

For the reading prompt, participants were given the following description about Ebola and asked to fill in a blank box by detailing the animals mentioned in the description:

> *The Ebola virus causes an acute, serious illness which is often fatal if untreated. Ebola virus disease (EVD) first appeared in 1976 in 2 simultaneous outbreaks, one in what is now, Nzara, South Sudan, and the other in Yambuku, Democratic Republic of Congo. The latter occurred in a village near the Ebola River, from which the disease takes its name. Ebola is introduced into the human population through close contact with the blood, secretions, organs, or other bodily fluids of infected animals such as [animal 1], [animal 2], [animal 3], and [animal 4].*

The animals listed in the description (animals 1-4) were experimentally manipulated between participants. Participants were randomly assigned to read either a CDC–inspired set of animals (fruit bats, gorillas, monkeys, and chimpanzees; *n* = 81) or a WHO-inspired set of animals (fruit bats, monkeys, forest antelope, and porcupines; *n* = 70).

Next participants completed the Ebola susceptibility questionnaire. For each question, participants were asked, “How likely is it that [animal] can get Ebola?” (1 = *Very Unlikely*, 7 = *Very Likely*). Animals included both mammals and birds: bats, monkeys, zebras, meerkats, anteaters, giraffes, gazelles, storks, flamingos, cranes, vultures, and parrots.

Next participants completed the bite reporting questionnaire. For the bite reporting questionnaire, participants were told to *“imagine that you are on a safari and get bitten by an animal, but the bite just barely breaks the skin”* when considering whether they would report a bite to a health professional. Each question asked them to rate (1 = *Very Unlikely*, 7 = *Very Likely*), *“how likely would you be to report being bitten by a [animal]?”*

Last, participants completed the meat safety questionnaire. For the meat safety questionnaire, participants were asked to rate (1 = *Very Unsafe*, 7 = *Very Safe*), “*how safe you think it is for people in general to eat meat from each animal”* and to “*consider only immediate health risks from disease transmission.”*

## Results

The results were consistent with predictions based on the premise diversity principle. Participants in the WHO (diverse) wording condition (fruit bats, monkeys, antelopes, and porcupines) rated individual mammals as more susceptible to Ebola [*t*(150) = 3.70, *p* < .001, *d* = 0.6; Fig 4A], were more likely to report mammal bites [*t*(150) = 2.83, *p* = .005, *d* = 0.46; Fig 4B], and perceived mammal meat as less safe [*t*(150) = 2.66, *p* = .009, *d* = 0.43; Fig 4C]. Because some of the animal premises (monkeys and bats) were included in both prompts, and thus the effect of condition may have a reduced effect on ratings of these animals, we used linear mixed effects models to test whether there were interactions between condition and animal for each of our ratings. There was a significant interaction for the effect of wording condition on susceptibility ratings [*F*(6,900) = 9.38, *p* <.001, *ηρ*^2^ = .06] whereby the effect of condition was not significant for bats and monkeys [*t*(150) = .1, *d* = 0.02] but was significant for animals that were not included in the instructions [*t*(150) = 3.88, *p* < 0.001, *d* = 0.63]. There were qualitatively identical interactions between wording condition and mammals for bite reporting and meat safety [Bite reporting: *F*(6,900) = 8.58, *p* < .001, *ηρ*^2^ =.05; Meat safety: *F*(6,900) = 5.74, *p* < .001, *ηρ*^2^ = .04].

**Fig 4.**
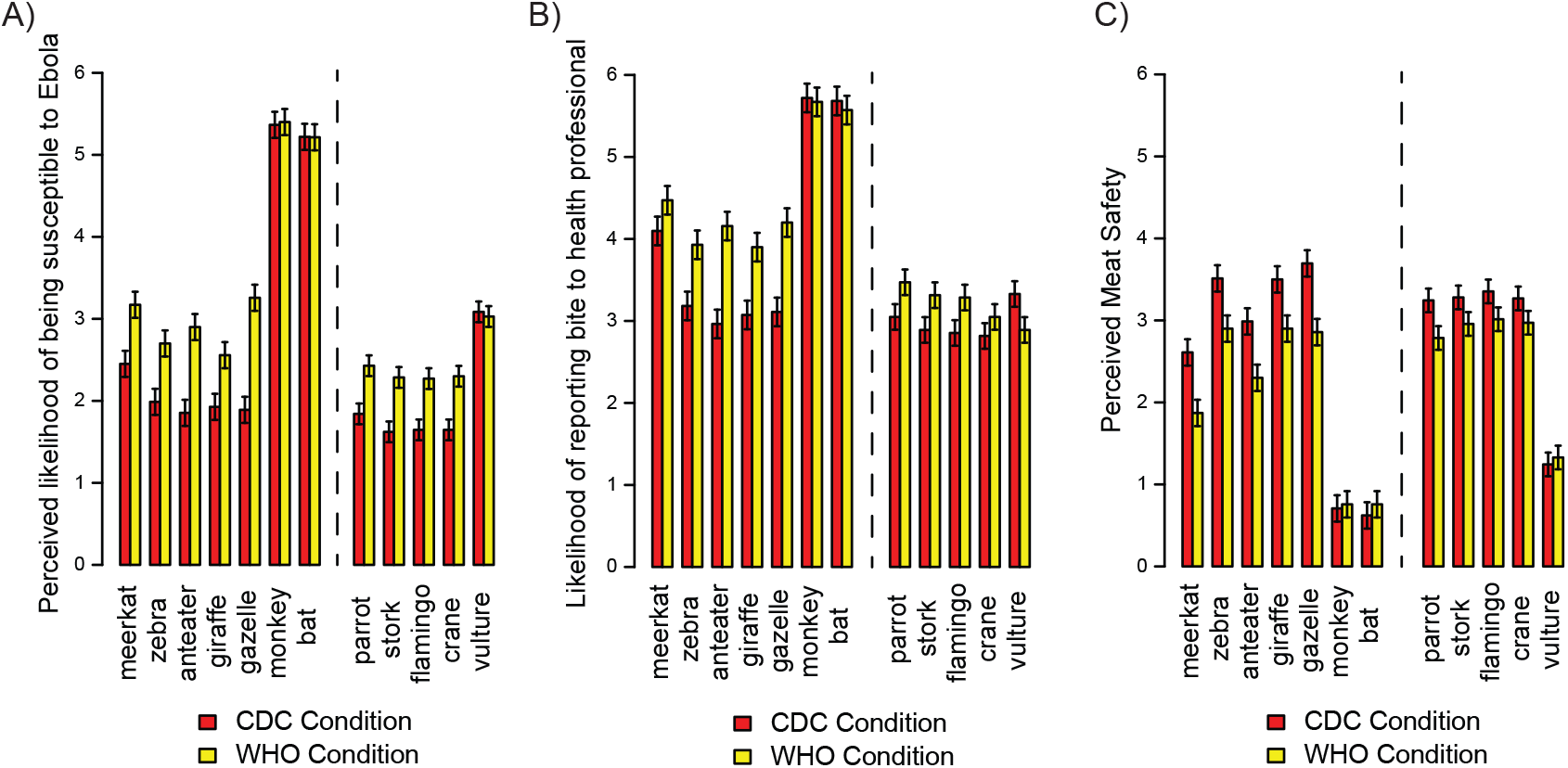
Study 2 results. (A) Perceived susceptibility of animals to Ebola. (B) Intentions to report animal bites (C) Perceived meat safety. Error bars reflect 95% within-subject confidence intervals.

The WHO (diverse) wording condition also increased perceptions of birds’ susceptibility to Ebola [*t*(150) = 2.06, *p* = .040, *d* = 0.33] but did not significantly increase intentions to report bird bites [*t*(150) = 1.10, *d* = 0.18] or lower perceptions of meat safety [*t*(150) = 1.28, *d* = 0.21]. Like ratings of mammal susceptibility to Ebola, there was a significant interaction [*F*(4,600) = 5.76, *p* < .001, *ηρ*^2^ =.04] whereby there was no effect of condition for vulture susceptibility ratings [*t*(150) = 0.19, *d* = 0.03], but there was an effect for other birds [*t*(150) = 2.55, *p* = .01, *d* = 0.42]. This was not expected a priori and may have to do with vultures’ tendency to eat carrion, making perceptions of their overall exposure to Ebola higher. This interaction was also present for bite reporting, *F*(4,600) = 4.06, *p* = .003, *ηρ*^2^ = .03, but did not reach significance for meat safety ratings, *F*(4,600) = 1.95, *p* = .10, *ηρ*^2^ = .01.

In addition to our primary tests of the effect of wording condition, we used linear regression to test whether the effect of wording condition on bite reporting and perceptions of meat safety was mediated by its effect on Ebola susceptibility ratings. First, we found that Ebola susceptibility were significantly associated with bite reporting and meat safety perceptions for both mammals and birds, even after taking into account the effect of wording condition [Mammal bites: standardized *b* = 0.47; *t*(149) = 6.40; *p* < .001; Mammal meat: standardized *b* = −0.43; *t*(149) = 5.67, *p* < .001; Bird bites: standardized *b* = 0.51; *t*(149) = 7.23, *p* < .001; Bird meat: standardized *b* = −0.45; *t*(149) = 6.01, *p* < .001)]. Next, we found that including Ebola susceptibility in the regression model with the effect of wording condition made the effect of condition non-significant for all models [Mammal bites: *b* = 0.18; *t*(149) = 1.20; Mammal meat: *b* = −.18; t(149) = -1.16; Bird bites: *b* = .009; *t*(149) = 0.063; Bird meat: *b* = −0.06; *t*(149) = 0.43], suggesting that the effects of condition on meat safety and bite reporting were fully mediated by the effect of the different wordings on participants’ perceptions of Ebola susceptibility. A bootstrapping procedure was used to test whether the indirect pathways between condition and bite reporting and condition and meat safety ratings through perceptions of Ebola susceptibility were significantly different from zero (Preacher & Hayes, 2008). None of the 95% bias-corrected confidence intervals included zero, suggesting that there were significant indirect pathways between wording condition and bite reporting and meat safety for both mammals and birds [Mammal bite reporting: 0.27, bias–corrected 95% CI (0.12, 0.49); Mammal meat safety: −0.25, bias-corrected 95% CI (-0.46, −0.11); Bird bite reporting: 0.17, bias-corrected 95% CI (0.003, 0.348); Bird meat safety: −0.15, bias-corrected 95% CI (−0.334, −0.012)]. Altogether, these analyses suggest that the effect of wording condition increased perceptions of disease transmission risk (through bites or wild game meat) by increasing the diversity of animals participants believe to be susceptible to Ebola.

## General Discussion

Results from both studies indicate an important role that cognitive research can play in combating emerging zoonoses. Although rarely studied within the One Health literature on zoonoses, humans’ inferences about risk are central to their interactions with potential disease vectors. We found that cognitive principles related to premise number and diversity impact individuals’ perceptions of zoonotic disease transmission risk and associated health behaviors. To the extent that people believe that it is possible for many diverse species to transmit diseases to one another, they become more wary of their own risk of infection.

An experiment based on CDC and WHO Ebola virus factsheets further revealed that individuals’ inductive reasoning strategies can be harnessed to make communications about disease risk more effective. Through the use of cognitive framing strategies, it may be possible to reduce adverse contact with animals and increase rapid reporting of potential disease exposure. Such interventions may be particularly effective for rural communities in remote areas that are difficult to reach with other interventions. These results have the potential to contribute to One Health goals of identifying low-cost strategies for reducing emerging disease risk before outbreaks occur [6, 7].

The present results are the first to suggest that inductive reasoning processes studied in cognitive psychology also influence health behaviors. With such connections established, future One Health studies on disease transmission risk perception would benefit from even stronger connections with cognitive research. One question is how the present results may generalize to expert populations, such as in preparing medical experts prior to work in the field. With experts, reasoning is often based on causal knowledge rather than heuristics like premise number and diversity [24–26]. Thus, communications based on such principles may be less useful for conveying disease risk to experts.

A second question is how people judge risks from different species. Here we focused on person-level characteristics that relate to perceived risk of animal contact (bites and game meat), averaging over differences between species. However, not all animals are associated with the same zoonosis risk, and it will be important to understand how to tailor communications to impact species selectively. For example, bats have a very strong association with emerging zoonosis [4, 27, 28], and in some cases it may be useful to tailor messages to focus on bats specifically. Although bats were associated with high levels of intended bite reporting and were perceived as being unsafe to eat, participants also may have underestimated the risks bats pose to other wildlife. Indeed, participants rated disease transmission risk between bats and other mammals as lower than for more typical mammals. Because wildlife-livestock interactions are a major driver of emerging zoonosis [29], this finding suggests that people may underestimate the risk of grazing wildlife near bat habitats.

The present research is primarily aimed at building interdisciplinary connections between One Health research and cognitive psychology. Still, the current results may have implications for basic psychological research on contagion as well. The law of contagion is a prominent social psychology construct that describes people’s tendencies to believe that negative (and positive) properties, including diseases and social ills, can be transmitted to objects or people through mere contact [18, 30]. Current theories of sympathetic magical thinking often make distinctions between the law of contagion and the law of similarity, a separate construct that describes the belief that objects that share surface features also share deeper common essences (e.g., leading to disgust with fudge shaped like dog feces, and beliefs that voodoo dolls can affect the person they resemble [31]). The present results suggest that the laws of contagion and similarity may not be fully separate, and similarity-based effects may influence perceptions of contagion. Indeed, theories suggest that inductive reasoning principles like premise number and diversity can increase generalization of properties (such as disease susceptibility) via similarity-relationships between known and novel/unknown examples. For example, the diverse prompts in our second experiment may have increased perceptions of Ebola susceptibility by increasing the likelihood that the unknown examples would match the known examples in some respect. A major question in cognitive psychology is how different respects [32] in which items can be similar (e.g., sharing the same category [11]; matching surface or internal properties [33,34]; or playing the same or complementary roles in a causal-relational system [35] impact generalization of novel/unknown properties. Although our data does not distinguish between these different candidate theories for similarity-based transfer of contagion, the results are suggestive that beliefs about contagion can be transferred via such similarity relationships.

The present results also inform basic research on induction. Social psychological research on health behaviors suggests that individual differences, such as sex [36], disgust sensitivity [20], and risk sensitivity [21], play a key role in people’s judgments about health risk. However, both health behaviors and individual differences in general have largely been ignored in cognitive psychological approaches to induction. To understand the influences of induction on real-world public health scenarios like zoonoses risk, it will be important to combine insights from more social domains with cognitive research. Indeed, in both of our studies, individual differences played a strong role in generalization. For example, in the second study, individual differences in beliefs about interspecies disease transmission mediated the effect of condition on bite reporting and meat safety judgments. For birds, these associations were significant even when the main effect of wording condition was not. These results suggest that, in domains like disease transmission, there are key individual difference factors in contagion beliefs that research on induction has not yet taken into account. Given that the paradigm we have developed reveals both psychometrically robust individual differences in health behavior intentions and strong effects of experimental manipulations, the present studies offer model paradigms that future experiments may use to further investigate person-by-situation interactions involving inductive reasoning principles such as premise number and diversity.

In conclusion, emerging diseases from animals pose a substantial public health threat, yet little is known about how people judge risks associated with different drivers of zoonoses. The present studies illustrate that basic cognitive principles related to inductive reasoning not only impact individuals’ perceptions of disease risk and associated health behaviors, but also can be harnessed for tailoring messages to properly convey risks associated with emerging zoonoses.

